# Glycine molecule radical: Predicted properties and dipeptide formation

**DOI:** 10.64898/2026.07.07.736934

**Authors:** Jaroslaw Synak, Jacek Blazewicz

## Abstract

The formation of the first peptide bonds under prebiotic conditions remains an open question. We used density functional theory calculations with the B3LYP functional to predict a novel pathway of peptide bond formation, which could have taken place without the sophisticated catalysts used by modern biological systems, utilising only radical chemistry. To make our investigation more extensive, the properties of intermediates (glycine-derived radicals) were thoroughly analysed, using the DFT, resonance hybrid model and orbital hybridisation. These methods shed more light on the exact nature of processes which should take place, explaining why this pathway could be favoured by the system. The result is a series of reactions, which without any sophisticated catalysts and with relatively low electronic energy barrier (<20 kcal/mol) can lead to formation of dipeptides, suggesting also a possible starting point for further peptide chain extension.

## Introduction

The transition from RNA World to protein-based life remains an unsolved problem in modern science. One of the main challenges is finding a plausible pathway, which would allow peptide bonds to form, without sophisticated biological machinery.

The RNA World hypothesis posits that, in the beginning, RNA molecules served both as information carriers and catalytic molecules. It is one of the most well-researched origin of life frameworks, with many arguments and results corroborating it ***Robertson and Joyce(2012);Saito(2022)***. Despite active research in the area, RNA World’s eventual transition into protein-based metabolism remains an open question. It is reasonable to assume that first peptides were made out of a limited set of aminoacids. Glycine is frequently considered one of the best candidates for such a primordial building block, due to its simple structure ***van der Gulik and Speijer(2015)***. Taking this into account, diglycine molecule formation becomes the simplest, but still plausible example of early peptide synthesis, hence it is the main focus of this article. It should be mentioned that the method proposed by the authors has a potential to be more general, with diglycine serving merely as a minimal test case.

Since the research into new peptide bond formation pathways poses a great challenge, it is immensely difficult to demonstrate complete and prebiotically plausible methods ***Frenkel-Pinter et al.(2022)***. Nevertheless, at least several possible pathways have been demonstrated empirically:

- the reaction is much more efficient, when thioester intermediates are used ***Frenkel-Pinter et al.(2022)***,
- peptides can form as chemisorption products of bifunctional amino acid vapours on the surface of silica and alumina***Basiuk et al.(1990)***,
- cysteine based catalyst could form in prebiotic conditions and catalyze peptide ligation ***Foden et al.(2020)***,
- chemoselective aminonitrile coupling in water (accepts all 20 basic aminoacids) ***Canavelli et al.(2019)***.

Simultaneously, considerable effort has been put into finding new chemical pathways of formation of simple aminoacids, mainly glycine, which would explain their presence in the environment. Many possible reaction routes, most using simple molecules as substrates, have been reported over the years, including mineral catalysed-synthesis, electric field enchancing effects and extraterrestial origin ***Carrascoza Mayén et al.(2019);Chimiak et al.(2024);Carrascoza et al.(2023)***; ***Mates-Torres and Rimola(2026);Cobb and Pudritz(2014)***.

As described in ***Szostak et al.(2017b)***, the emergence of life on Earth can be intuitively separated into four distinct phases:

- α- inorganic compounds,
- β- nucleotides,
- γ- polynucleotide polymers,
- δ- self-organized polynucleotide chains.

There has been a lot of effort put into investigation of γ to δ transition ***Szostak et al.(2016***, 2017a);***Synak et al.(2020***, 2022, 2024) with foundations laid by Takeuchi and Hogeweg ***Takeuchi and Hogeweg*** (2009,2012). This article explores further evolution, going beyond theδphase, investigating one of potential paths of peptide synthesis in a relatively simple environment. It employs computational methods (mainly Density Functional Theory) to predict a prebiotically plausible reaction pathway, leading to peptide bond formation, which utilises radical chemistry, with no external catalysts necessary. This process could signify the first step towards protein-based biology.

## Materials and methods

The authors used Gaussian 16 ***Frisch et al.(2016)*** to perform all essential calculations, especially its IRC (Intrinsic Reaction Component) functionality. The method utilised was DFT with B3LYP functional with unrestricted electrons and the base 6-31G(d), as it provides sufficient precision in many contexts ***Tirado-Rives and Jorgensen(2008)*** with a reasonable computational time, allowing the authors to explore many different scenarios and pathways.

Both the necessary infrastructure and the license were provided Poznan Supercomputing and Networking Center, which shared its cluster called Eagle (its total computational power reaches 1.7 PFlops) ***PCS(2026)***.

Each of the reaction analysed in the article has been found with a combination of theoretical and computational methods. First the reagents were examined using mainly resonance hybrid model and organic chemistry basics to identify which interactions were most reasonable. The second stage was to calculate the total energy of reagents and the products to determine if the change in energy would be reasonable. The third and simultaneously the most complicated stage was guessing the transition state and using Gaussian “relaxed scan” function to find the best distances between atoms and angles between them. This examination could be conducted extensively, since the molecules researched in this work are relatively small.

## Results

In this section, all results of the authors’ considerations and calculations are presented. First, the exact way a diglycine peptide bond can form is described and energy of this reaction is discussed. The next subsections covers the reactions leading to the most important component of the afore-mentioned process - glycine acyl radical and a molecule, which was not necessary for this reaction, but the authors decided to describe it nonetheless - “simple” glycine radical. Their properties, how they emerge and theoretical models explaining their eletron structure will be covered.

### Peptide bond formation

The main staple of this work is to describe the results of the calculations, regarding the reaction between the glycine acyl radical and the glycine molecules. The electronic energy barrier of this reaction is lowered, first and foremost due to the partial positive charge of the acyl carbon atom and partial negative charge of the nitrogen atom in the glycine molecule (due to its high electronegativity), causing the atoms to slightly attract themselves as a result of pure electrostatic force.

The reaction progress was thoroughly calculated using Gaussian with the settings described in the Materials and methods section. The electronic energy barrier predicted by the tool was equal to19.34 kcal/mol. The process is endergonic, which is common for peptide bond formation, with the products having8.13 kcal/mol more energy than the reactants. The results of IRC calculation are presented in the Fig.1, which presents how the energy of the system changes during the entire process.

The mechanism is surprisingly simple. The acyl group approaches the amino group of the glycine molecule and a peptide bond is formed almost immediately. Simultaneously one of the hydrogen atoms of the amino group is rejected.

**Figure 1.**
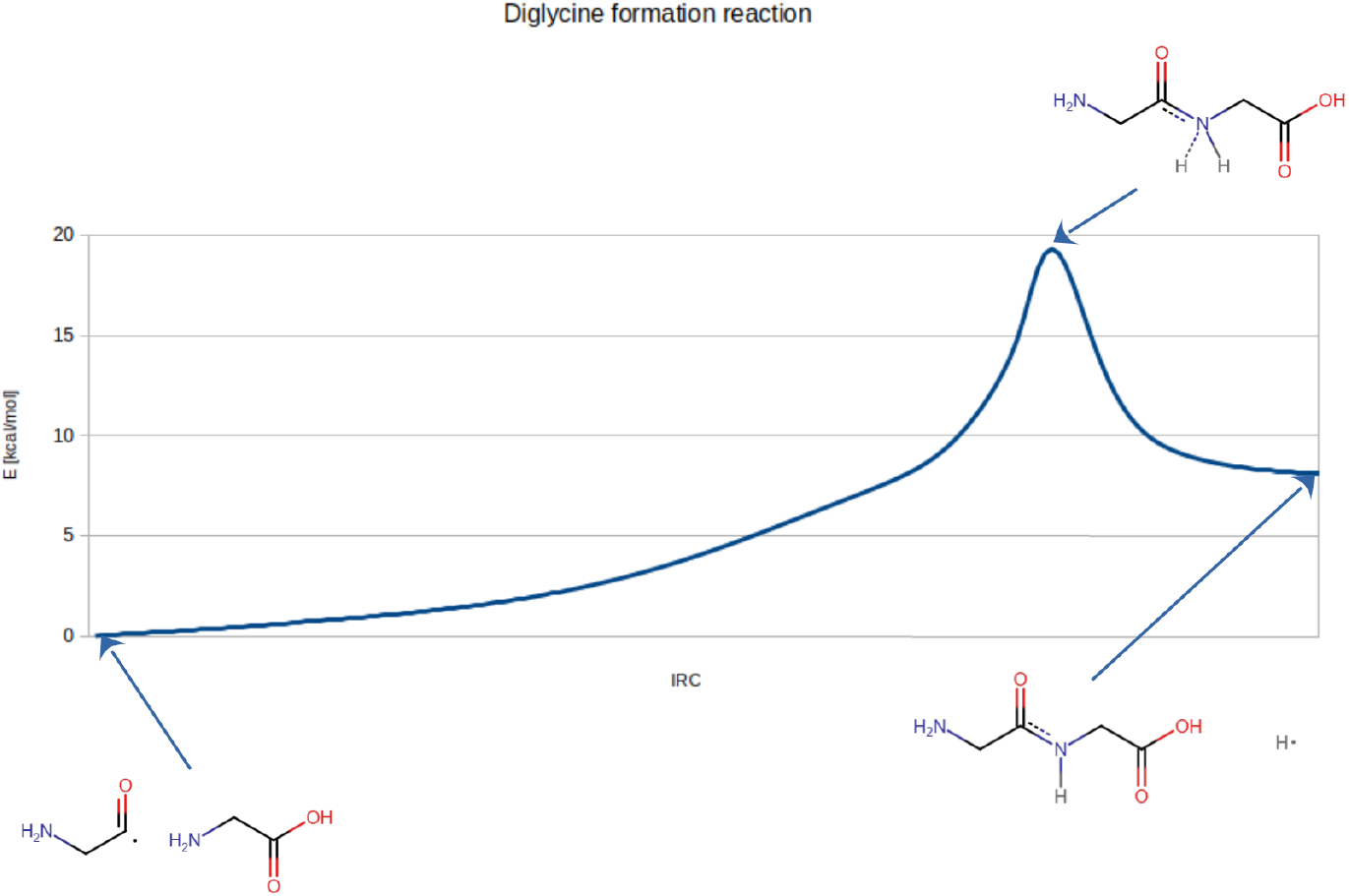
**The changes in energy during the peptide bond formation, leading to diglycine. The process is endergonic, which is not unusual for peptide formation. Small size of the hydrogen atom allows it to escape virtually instantly, ensuring that the products are favoured.**

### Simple glycine radical

This radical forms, when one of the CH bonds in glycine is broken. The authors analysed its properties using the resonance hybrid model and identified three main contributing structures (Fig2).

**Figure 2.**
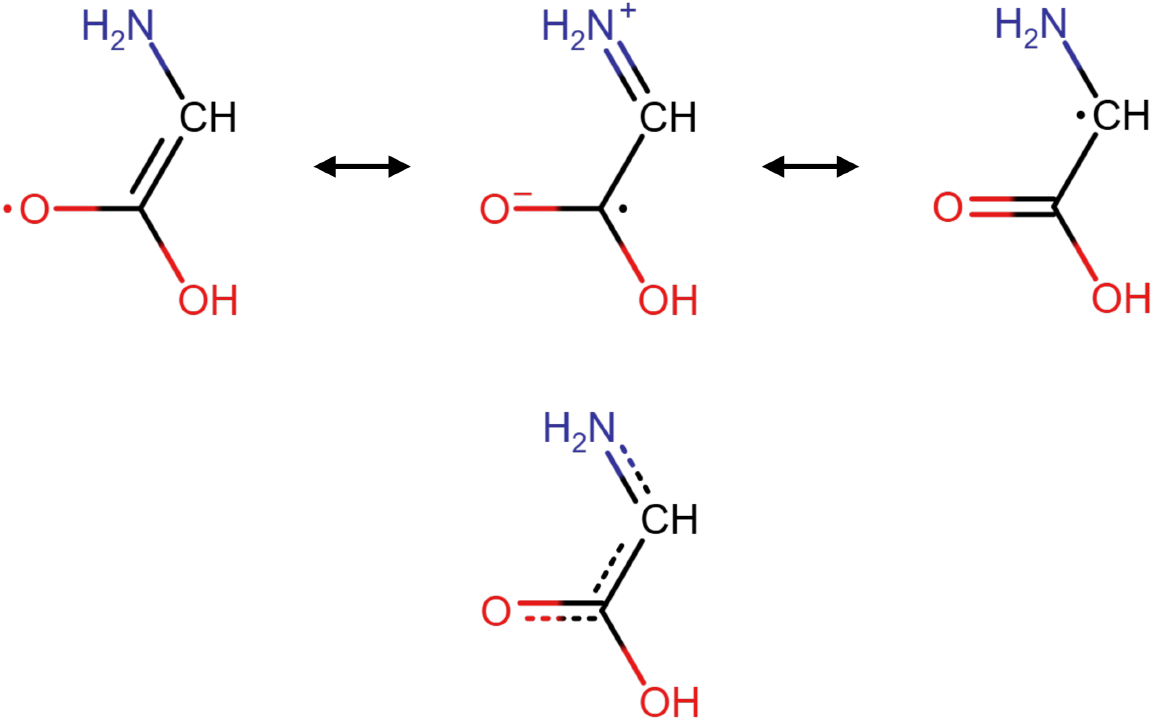
**Simple glycine radical as a resonance hybrid. Three contributing structures and the resonance hybrid below.**

The resonance hybrid model was corroborated by calculations regarding the Mulliken charge of atoms. The nitrogen atom should be charged more positively than in glycine, while the oxygen atom with a double bond should have a slightly more negative charge. While the exact numbers were strongly dependent on the isomer/conformer taken into account, the difference for nitrogen atom fell into the range of0.03 − 0.08*a.u*.and the difference for the oxygen atom was similar:0.04 − 0.07*a.u*..

It is important to note that the first contributing structure stiffens the CC bond, which becomes something between a single bond and a double bond. The same thing happens with CN bond in the molecule (the middle contributing structure), which results in a flat geometry, confirmed by Gaussian. Both C atoms and the nitrogen lean strongly towards sp2 hybridisation, the fact which is further supported by angles between bonds of each of these atoms, which nears120 ^°^ (112.28 − 125.89).

Due to this mixed nature of CC bond, the rotation around it is much more difficult, which results in cis-trans isomerism (Fig.3). This fact is proved by the energy required to switch from the Z-isomer to the E-isomer by rotation -18.70 kcal/mol.

**Figure 3.**
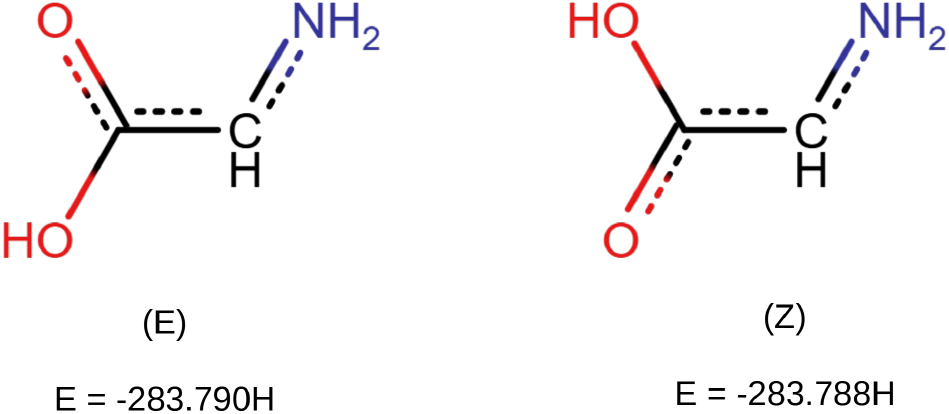
**Simple glycine radical - EZ isomerism.**

#### Synthesis

The simplest reaction leading to the radical was to break one of CH bonds, using atomic hydrogen (Fig.4). The electronic energy barrier for this particular process is predicted to be only 3.19 kcal/mol. It is a favourable reaction, releasing around26.49 kcal/mol and leading to the (E)-isomer. Due to the resonance, the glycine radical is quite stable, so any sufficiently active radical should be able to break the bond.

**Figure 4.**
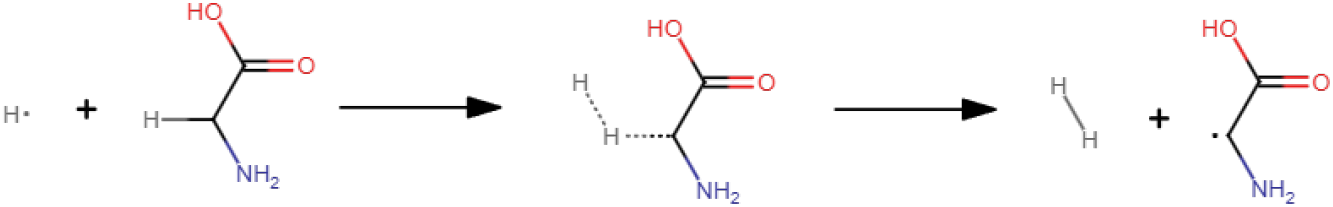
**The reaction of glycine and atomic hydrogen. CH bond is broken and a hydrogen molecule is formed.**

There is an alternative path with a slightly higher electronic energy barrier (4.47 kcal/mol) and releasing less energy (17.41 kcal/mol), resulting in the slightly less stable (Z)-isomer.

#### Alternative radicals

Since glycine molecules have three groups of hydrogen atoms (OH, CH and NH), two alternative radicals have been investigated. One emerges when the hydrogen atom in the COOH group is removed, however this variant is unstable and quickly decays into a methylamine radical and carbon dioxide in a process vastly similar to the one taking place during Kolbe electrolyisis ***Vijh and Conway(1967)*** (Fig.5).

**Figure 5.**
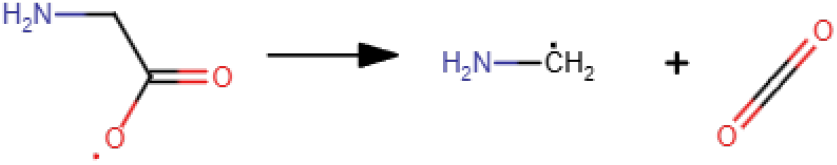
**The glycine molecule decay after losing its oxygen-bonded hydrogen atom.**

The other possibility is to break one of NH bonds. In that case, the radical is significantly less unstable with the hydrogen from the OH group forming a hydrogen bond with the nitrogen atom. However, since the hydrogen atom comes close to the nitrogen atom, only11.75 kcal/mol for the OH bond to be broken and an full NH bond to form. This results in the same decay process, which has been mentioned above (fig 6) with the reaction being exergonic (its total energy balance is equal to14.89 kcal/mol).

**Figure 6.**
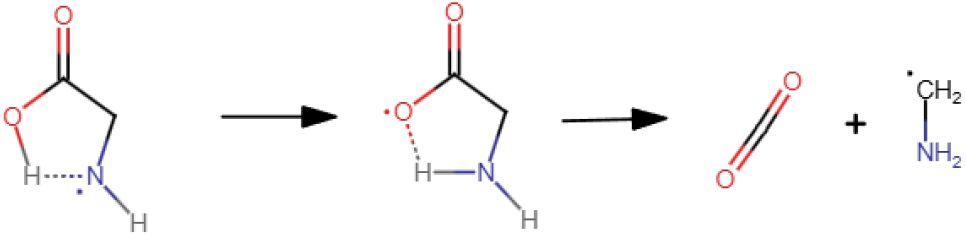
**The glycine molecule decay after losing one of its nitrogen-bonded hydrogen atoms.**

### Acyl radical

This subsection covers the properties of the most important reactant of the protein bond formation reaction - the glycine acyl radical molecule. Its structure is shown in the Fig.7.

**Figure 7.**
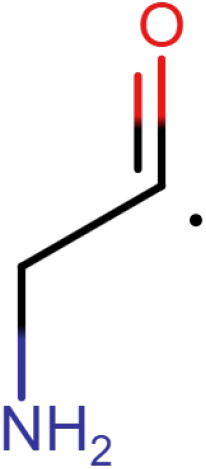
**The glycine acyl radical structure.**

#### Synthesis

This radical, according to predictions, can form in a two step process. First, a glycine molecule reacts with a hydrogen atom, losing the amino group and releasing a lot of energy in the process (31.54 kcal/mol). The activation energy is equal to approximately12.92 kcal/mol. Due to higher electronic energy barrier, this reaction is much slower than the simple radical formation, but is still possible. The reaction is shown in the Fig. 8

**Figure 8.**
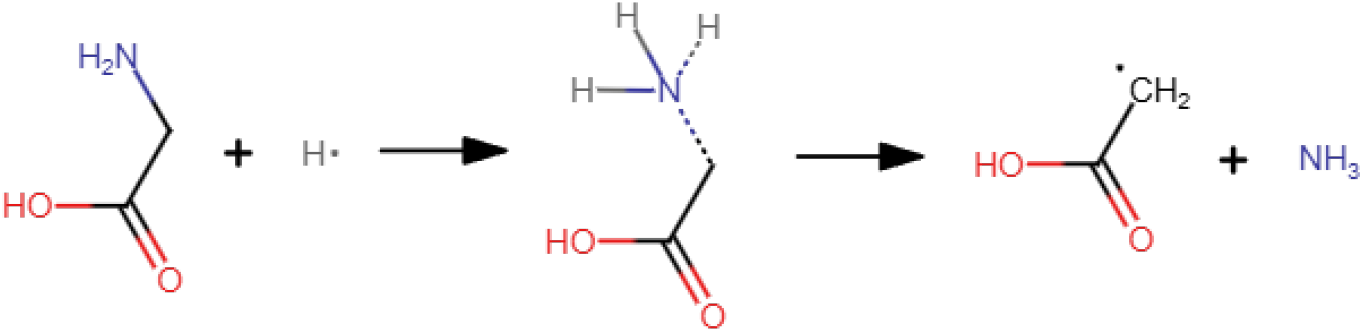
**Glycine molecule losing the amino group reaction.**

If the energy during this reaction is not dissipated, the products can step into another reaction. The ammonia molecule can reattach, temporary forming the glycine molecule again, while the hydrogen atom being released, reacts with the OH group, detaching it from the carbon atom and forming a water molecule. The reaction is presented in detail in the Fig.9. The electronic energy barrier for this process is high, circa45.21 kcal/mol, but it is only slightly higher than that of the previous reaction if the initial glycine molecule and the hydrogen atom are took as the reference point (13.67 kcal/mol). The corresponding values of the energy balance are18.83 kcal/mol and −12.71 kcal/mol.

**Figure 9.**
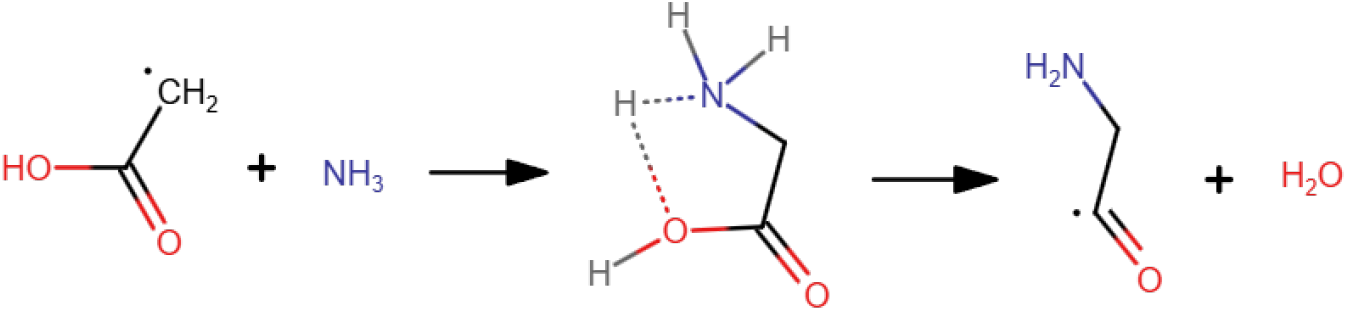
**Ammonia reattaches, forming glycine and the free hydrogen atom reacts with the OH group.**

The changes of energy of the two step process described above are presented in the Fig.10.

**Figure 10.**
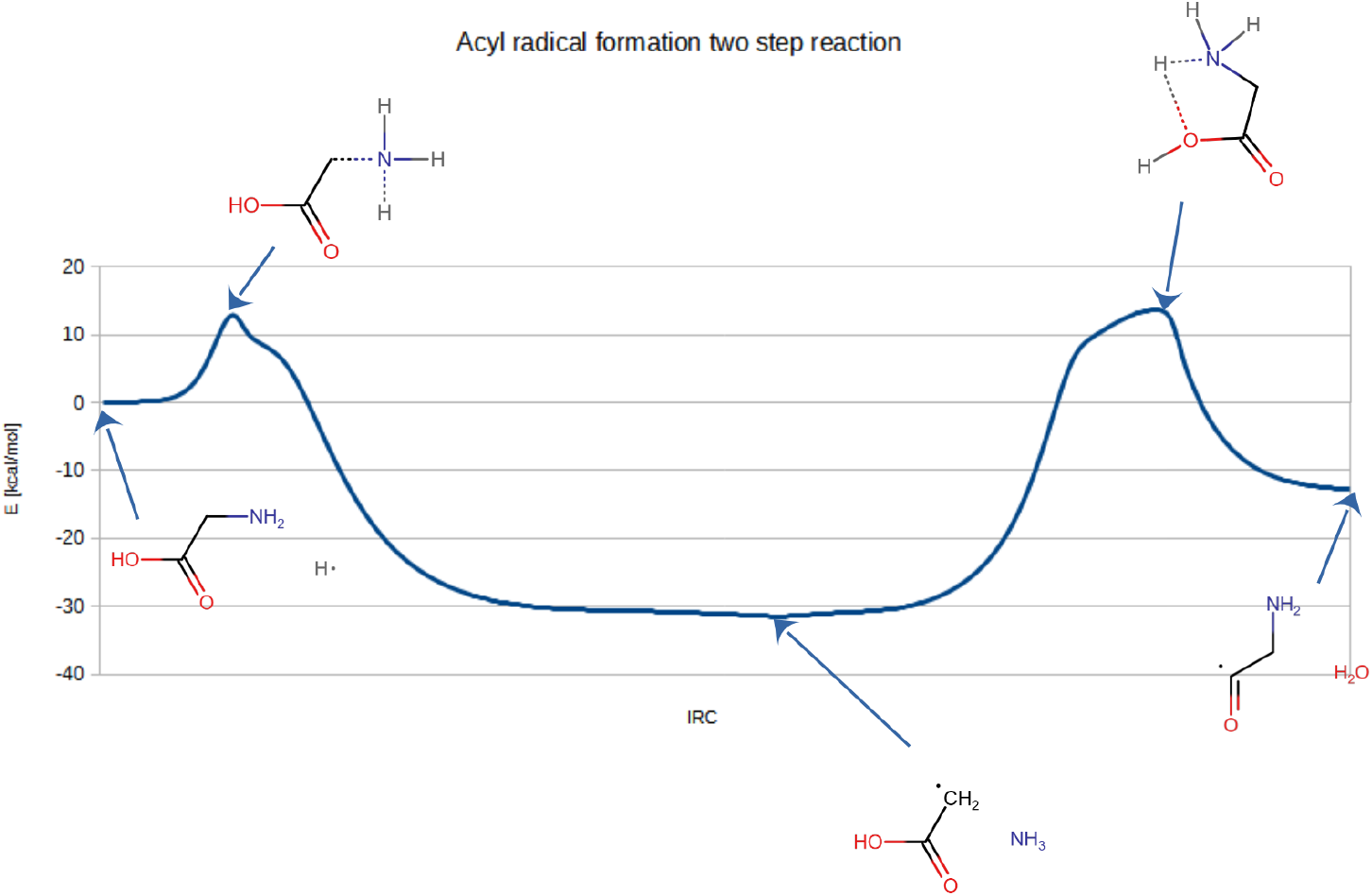
**Two step process leading to the glycine acyl radical formation.**

It should be mentioned than the transitive radical of acetic acid is stabilised by resonance, that partially explains its low energy (Fig.11).

**Figure 11.**
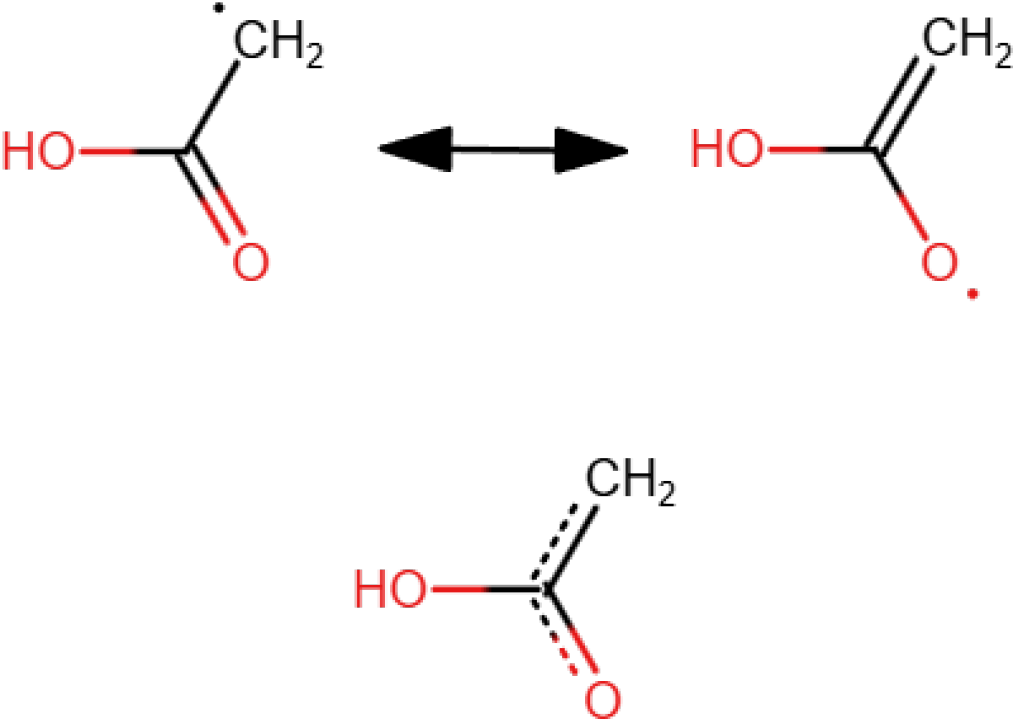
**Acetic acid radical is partially stabilised by resonance. Two contributing structure and the resonance hybrid below.**

The reaction presented may seem overcomplicated, however the direct reaction of atomic hydrogen with the OH group needs high activation energy -29.29 kcal/mol.

## Discussion

The methods of theoretical chemistry and computer simulations are an important tool of research, alongside wet experiments. It should be noted that using machine learning methods to accelerate calculations and reduce errors shows great promise ***Keith et al.(2021)***.

The method used to obtain the results (B3LYP) is known to have severe limitations for larger systems, especially for multiple (more than 4) C-C bonds ***Check and Gilbert(2005)***. Understanding this is important, as there is no universal model, which works for every scenario. The molecules analysed in this paper are shorter, hence errors should be in an acceptable range.

The subject of the publication - the properties of glycine molecule radicals, is to complement current investigation into glycine molecule properties and its potential role in the primordial soup. Approaching the problem from the point of view of quantum chemistry is a well-estaablished field of research ***Rimola et al.(2022)***. One direction is to explain the emergence of glycine and other important building blocks of life in space, as it is evident that organic compounds can occur outside of Earth ***Kwok(2016)***. Various routes of glycine synthesis in interstellar medium have been proposed, some involving reactive radicals in a gas phase ***Pilling et al.(2011);Woon(2002)***, others taking place in a solid phase, for instance interstellar ice ***Rimola et al.(2010);Carrascoza et al.(2023)***.

## Conclusion

In this paper, using Gaussian and theoretical chemistry, glycine radical and its acyl radical were analysed. The most important result is a possible formation of peptide bond, the electronic energy barrier of which remains in an acceptable energy range (19.34 kcal/mol). Since the presence of radicals has to be corroborated, a possible synthesis method has been searched for and described in this publication with prediction of electronic energy barrier and energy balance for every reaction along the way.

The results described above should be a good starting point for anyone researching these radical pathways in the future. Due to limitations posed by imperfections of our current quantum chemistry models, energy values, being the main staple of this article, are only approximations. Nonetheless, the models point in an interesting direction, showing a possible peptide bond formation, which utilises no complex catalysts, relying solely on radical chemistry. Glycine radical molecule has been thorougly analyzed as a possible candidate, which could in theory lead to peptide bond formation, but the simulations shown, the acyl radical was a much better candidate. However, the former molecule was still included in the paper, both to discuss its interesting properties and serve as a guide for further research regarding in role in various prebiotic pathways.

## Supporting information

Gaussian calculation logs

## Acknowledgments

This work was partially supported by statutory funds from the Institute of Computing Science of Poznan University of Technology, grant number: 0311/SBAD/0767. The computations were carried out using the infrastructure and Gaussian license shared by Poznan Supercomputing and Networking Center, Jana Pawla II 10; 61-139 Poznan, Poland (grant pl0295-04).

